# Elucidating the Characteristics and Clonal Evolutionary Trajectory of Influenza Neuraminidase Broadly Reactive B Cell

**DOI:** 10.1101/2025.02.23.639775

**Authors:** Chaohui Lin, Qian Yang, Lianyu Huang, Xin Wang, Zeli Zhang

## Abstract

Influenza virus neuraminidase (NA) is receiving increasing attention as a target for universal flu vaccines. Several broad NA inhibition monoclonal antibodies (BImAbs) targeting the highly conserved enzymatic pocket have been previously described. However, the molecular characteristics, clonal evolutionary trajectory, and B cell sources of BImAbs remain poorly understood. Here, using NA-mutant probes, we comprehensively profiled the immune signatures of NA-specific memory B cells (MBCs) from a healthy individual with NA cross-inhibition activity. From the NA-specific MBC repertoires, we identified a series of NA BImAbs with molecular features characterized by long HCDR3 regions with an “xxxDRxxx” motif, which exhibited broad inhibition against diverse influenza NAs. Clonal lineage tracing revealed that these BImAbs followed a clonal evolutionary trajectory encompassing classical MBC (cMBC) and atypical MBC (aMBC). Both cMBC- and aMBC-derived NA BImAbs displayed similar inhibitory activity against the influenza NAs. These findings enhance our understanding of the development of NA BImAbs and provide a foundation for the rational design of NA-based universal flu vaccines.

## Introduction

Influenza virus poses a significant public health threat, causing millions of cases, hospitalizations, and numerous fatalities annually (1, 2). Influenza A virus (IAV, subtypes H1N1 and H3N2) and influenza B virus (IBV, lineages Victoria and Yamagata) are responsible for seasonal outbreaks in humans. IAV, particularly H1N1, occasionally causes pandemics (3). Additionally, the spillover of avian influenza viruses (such as H3N8, H10N3, and H5N1) to humans has been reported (4-8). The recent spread of highly pathogenic avian influenza (HPAI) H5N1 among cattle and its spillover to humans has raised concerns about the potential pandemic risk (6-8). Influenza viruses undergo rapid mutations, which may enable them to escape the immune response induced by previous infection or vaccination. Therefore, developing a universal flu vaccine that provides broad and long-lasting immunity is essential and has been the focus of many researchers (9).

Influenza viruses possess two main glycoproteins: hemagglutinin (HA) and neuraminidase (NA), both of which are essential for viral replication. Compared to the highly variable head of HA, NA exhibits lower mutation rates and contains a relatively conserved enzymatic pocket targeted by the widely used antiviral drug oseltamivir (10, 11). Recent studies have characterized NA-specific antibody levels in humans, suggesting that these antibodies are associated with protection against the virus and a reduction in viral shedding (12-15). Additionally, several NA broad inhibition monoclonal antibodies (BImAbs) have been identified from human plasmablasts through unbiased screening or from pre-existing memory B cells (MBCs) using NA-mutant probes, providing broad cross-reactivity against diverse influenza viruses *in vitro* and *in vivo* (12, 16-18). Elucidating the conserved protective epitopes on NA targeted by these BImAbs has provided essential insights for the development of NA-based influenza vaccines (19).

Human MBCs represent a crucial component, providing long-term protection against the influenza virus, particularly its variants (20, 21). Recent studies have illuminated the significant phenotypic and functional heterogeneity within the human MBC compartment (21, 22). Compared with classical MBCs (CD27^+^, cMBCs), atypical MBCs (CD27^-^CD21^-^ FCRL5^+^CD11c^+^, aMBCs) have been identified in conditions such as malaria, HIV infection, and autoimmune diseases (23-26). In HIV-1 infected individuals, aMBCs exhibit a dual characteristic of diminished BCR signaling responsiveness but enhanced sensitivity to the Toll-like receptor pathway (27). In contrast, malaria-specific aMBCs can encode broadly neutralizing antibodies and are associated with protection (24, 28). Additionally, although the proportion of aMBCs in the peripheral blood of healthy individuals is low, their numbers can significantly expand following vaccination or infection (28, 29). Furthermore, influenza HA^+^T-bet^+^FCRL5^+^ aMBCs exhibit effector-like memory cell characteristics and can rapidly differentiate into antibody-secreting cells *in vitro* (30). However, whether aMBCs contribute to the NA broad-reactive MBC precursors and their clonal evolutionary trajectory remains unclear. Understanding the molecular features of NA broadly reactive MBCs, their working mechanisms, and developmental trajectories is crucial for the rational design of NA-based universal flu vaccines.

In the current study, we identified a healthy individual with NA cross-inhibition activity. By combining NA-mutant probe-based MBC sorting with single B cell BCR and transcriptome sequencing, we comprehensively profiled the immune phenotype of the NA-positive MBCs and discovered that a long heavy chain complementarity-determining region 3 (HCDR3) with an “xxxDRxxx” motif is an essential characteristic of NA broad-reactive B cells. Several isolated monoclonal antibodies with this feature exhibited broad inhibition activity against diverse influenza NAs. Clonal lineage tracing revealed that these B cell derived BImAbs followed a clonal evolutionary trajectory encompassing both cMBCs and aMBCs. Furthermore, we demonstrated that both cMBC- and aMBC-derived BImAbs displayed similar inhibition activity against the influenza virus *in vitro*.

## Results

### Identification of an individual with NA cross-reactive antibodies

NA inhibition (NI) antibodies against seasonal influenza strains have been detected in healthy adults (12, 31). To determine whether NA cross-inhibition antibodies could be identified, we tested the NI titers of 15 serum samples obtained from healthy adults in 2023-2024 against various NAs (Figure 1A-C). All samples exhibited detectable NI titers against NA from seasonal influenza strains, including N1 from A/Michigan/45/2015-H1N1, N2 from A/Kansas/14/2017-H3N2, and NB from B/Colorado/06/2017-Victoria (Figure 1C). While most NI titers against N3 (A/mallard/Minnesota/AI09-3100/2009-H6N3) and N5 (A/mallard/Sweden/86/2003-H12N5) were below the limit of detection, one serum sample (donor 3) demonstrated high inhibition against both N3 and N5 (Figure 1C). To further confirm the NA cross-reactivity, we purified total IgG from the serum of donor 3 and measured its binding activity against diverse NAs. The results indicated that the purified total IgG maintained reactivity against all tested NAs (Figure 1D-1E). These findings suggest that donor 3 possessed broadly reactive NA antibodies and likely maintained long-lived, broadly reactive MBCs.

**Figure 1.**
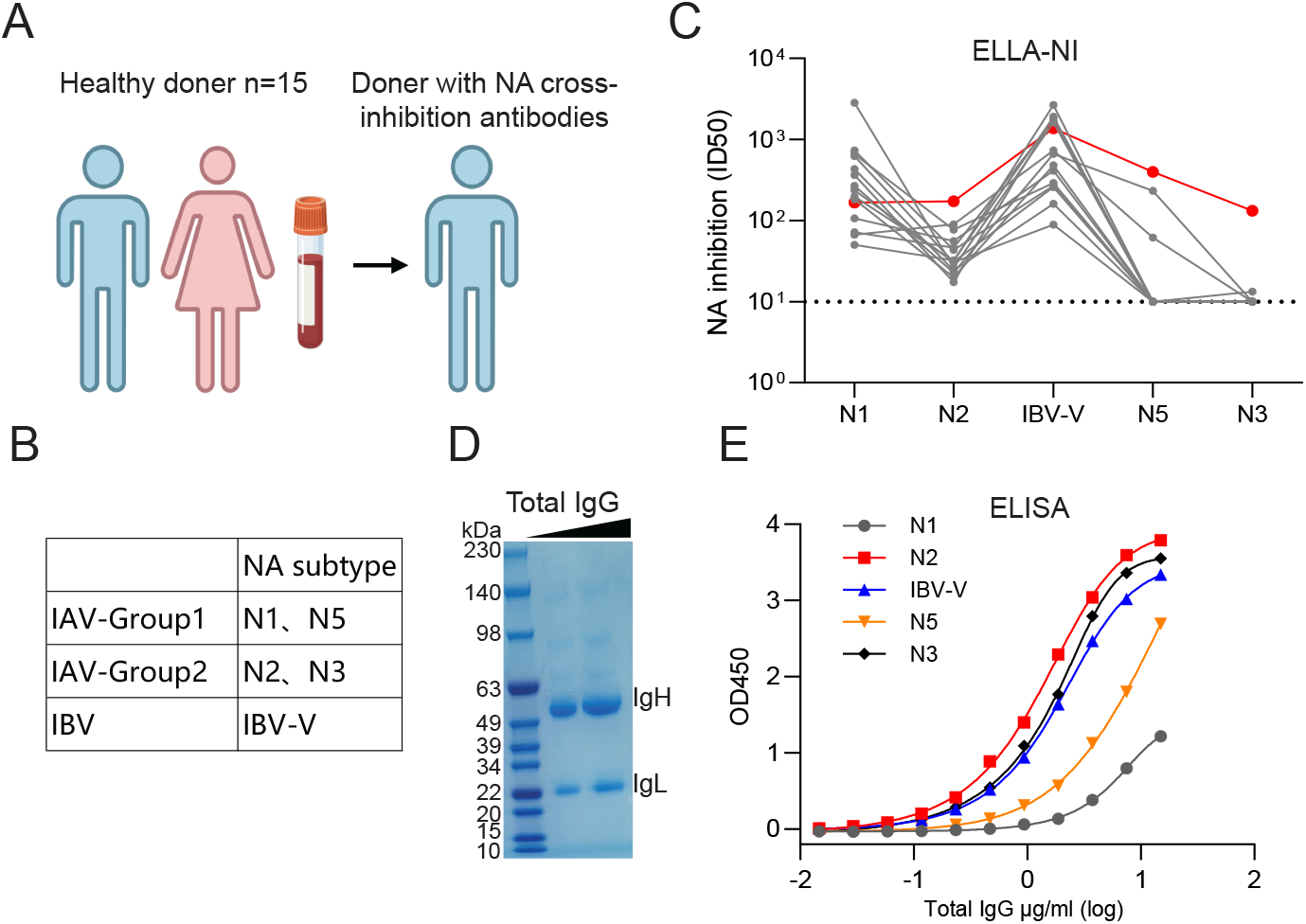
Identification of a donor with NA cross-reactive antibodies. (A) Schematic illustration of screening donors with NA cross-inhibitory antibodies from healthy adults. (B) IAV and IBV NA subtypes are utilized for NA-inhibition antibody screening in an ELLA assay. (C) The pre-existing NA-inhibition antibody titers in different donors. (D) Visualization of purified total IgG from donor 3’s serum by SDS-PAGE and Coomassie staining (E) Binding of purified total IgG to N1, N2, IBV-NB, N5, and N3 measured by ELISA. The recombinant N1, N2, IBV-NB, N5, and N3 used in Figures 1C and 1E were derived from strains A/Michigan/45/2015-H1N1, A/Kansas/14/2017-H3N2, B/Colorado/6/2017(Victoria lineage), A/mallard/Sweden/86/2003-H12N5, and A/mallard/Minnesota/AI09-3100/2009-H6N3, respectively.

### Single-cell RNA and VDJ sequencing profile the immune features of NA-specific MBCs in donor 3

Our recent study developed NA-mutant (NA-M5) probes that can specifically identify NA-specific MBCs (12). To investigate the potential broad NA-reactive B cell response in donor 3, we sorted 11,427 N2-specific MBCs from total MBCs (gated on live/CD14^−^/CD16^−^/CD3^−^/CD19^+^/IgD^−^/CD27^−^/^+^) using the N2-M5 probes (Figure 2A). Additionally, we collected non-N2-specific MBCs, stained them with NB-M5 probes, and enriched approximately 3,631 NB-specific MBCs (Figure 2A). Subsequently, the sorted N2 and NB-specific MBCs were pooled and subjected to 10X Chromium for single-cell RNA and VDJ sequencing. Based on their gene expression profiles, the NA (N2 and NB)-specific MBCs were classified into 15 clusters, reinforcing our prior observations that NA-specific MBCs are heterogeneous (Figure 2B) (12). Using SingleR algorithm annotation, NA-specific MBCs were further categorized into naïve-like MBCs (clusters 3, 4, 8, 9, and 10, black circle), class-unswitched MBCs (clusters 0, 1, 5, 6, 11, 13, and 14, blue circle), and class-switched MBCs (clusters 2, 12, and 7, red circle) (Figure 2B).

**Figure 2.**
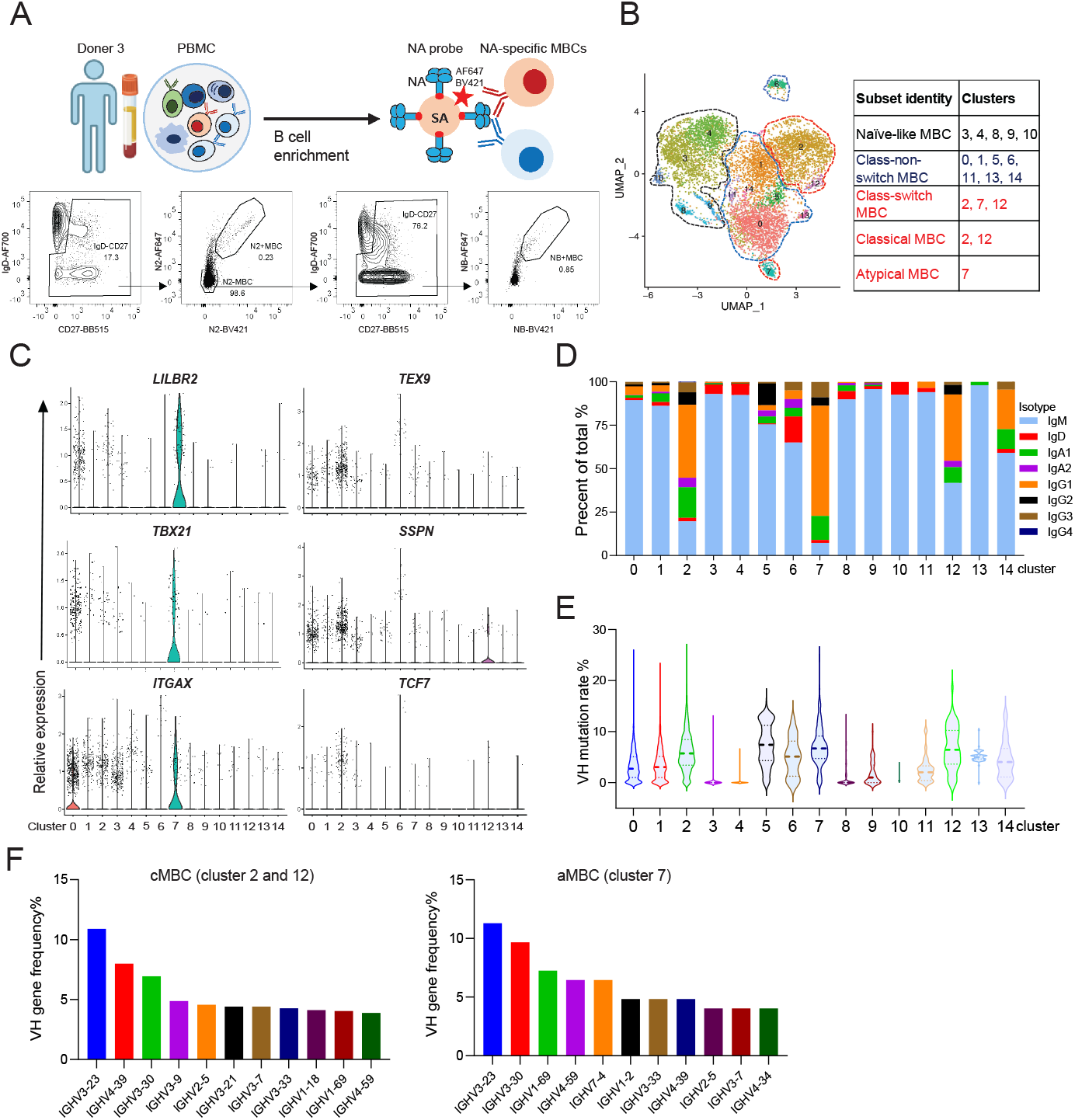
Immune profiling of the pre-existing NA MBCs in donor 3. (A) Schematic depicting our strategy for sorting NA-specific MBCs from enriched human B cells. (B) Integrated transcriptional uniform manifold approximation and projection (UMAP) analysis of distinct NA-specific MBC clusters. (C) The relative expression of marker genes across NA-specific MBC clusters. (D) Quantitative visualization of Ig subtypes used by NA-specific MBCs. (E) The VH mutation rates of NA-specific MBCs from different clusters. (F) The frequencies of top 10 VH genes usage by NA classical and atypical MBCs, respectively.

Among the NA-specific class-switched MBC population, two transcriptomically distinct subpopulations were defined as cMBCs (clusters 2 and 12) and aMBCs (cluster 7) using previously described MBC gene signatures (22, 32) (Figure 2B). As shown in Figure 2C, cMBCs exhibited high expression of *TEX9, TCF7*, and *SSPN*. aMBCs expressed higher levels of *LILRB2 (CD85d), TBX21 (T-bet)*, and *ITGAX (CD11c)*. To further discern the identity of each NA-specific MBC cluster, we analyzed the Ig repertoires and the somatic hypermutation (SHM) rates of the V gene of the heavy chain. Clusters 0, 1, 3, 4, 5, 6, 8, 9, 10, 11, 13, and 14 were dominated by the IgM isotype (Figure 2D), while clusters 2, 7, and 12 contained more IgA and IgG isotypes, consistent with the “class-switched MBC” classification by the SingleR algorithm (Figure 2D). Additionally, clusters 2, 12, and 7 had relatively higher VH mutation rates compared to other clusters (Figure 2E). Regarding Ig switch and SHMs, no significant difference was observed between the cMBC (clusters 2 and 12) and aMBC (cluster 7) compartments. We subsequently analyzed the top 10 VH genes utilized by cMBCs and aMBCs (Figure 2F). VH3-23 was dominant in both cMBCs and aMBCs with a frequency of approximately 11% (Figure 2F). Notably, both VH4-39/VH4-59 (used by N1 or N2-specific inhibition mAbs) and VH3-30/VH1-69 (utilized by NA BImAbs) were observed (12, 17, 33, 34), suggesting both NA strain-specific and broad B cell responses.

### Identification of several distinct BImAb clonotypes featuring an HCDR3 “xxxDRxxx” motif

Pan-NA BImAbs exhibit conserved structural signatures, including long HCDR3 loops (>18 amino acids) that enable deep penetration into the NA active site, coupled with elevated SHMs (12, 16, 17). Based on these features, we recently identified NA BImAb 4N2C4, which utilizes an Asp-Arg (DR) motif within its HCDR3 to mimic sialic acid interactions with conserved catalytic residues (12). This observation led us to hypothesize that the “DR” motif in the middle of a long HCDR3 loop constitutes a defining feature of NA BImAbs (Figure 3A-B). To validate this hypothesis, we used the following criteria to screen potential BImAbs from the BCR repertoires of donor 3: (1) HCDR3 > 18 amino acids, (2) the middle of the HCDR3 contains a “DR” motif, (3) VH mutation rate reaches approximately 10%, and (4) Ig isotype is IgG. We selected four clonotypes that adhered to these criteria (Figure 3A). Additionally, we chose clonotype C17, which contained signatures (1), (3), and (4), but lacked the “DR” motif in the HCDR3 and hypothesized that C17 was not a BImAb. The naming of these mAbs is based on the clonotype and the range of VH mutation rates. For example, C7-1: “C7” indicates “clonotype 7,” and “1” denotes “the highest VH mutation rate among its clonotype.” The VH and VL usage, mutation rate of the V gene, and CDR3 sequences are shown in Figure 3A. Furthermore, the recombinant IgG1 of these selected mAbs were expressed for functional characterization.

**Figure 3.**
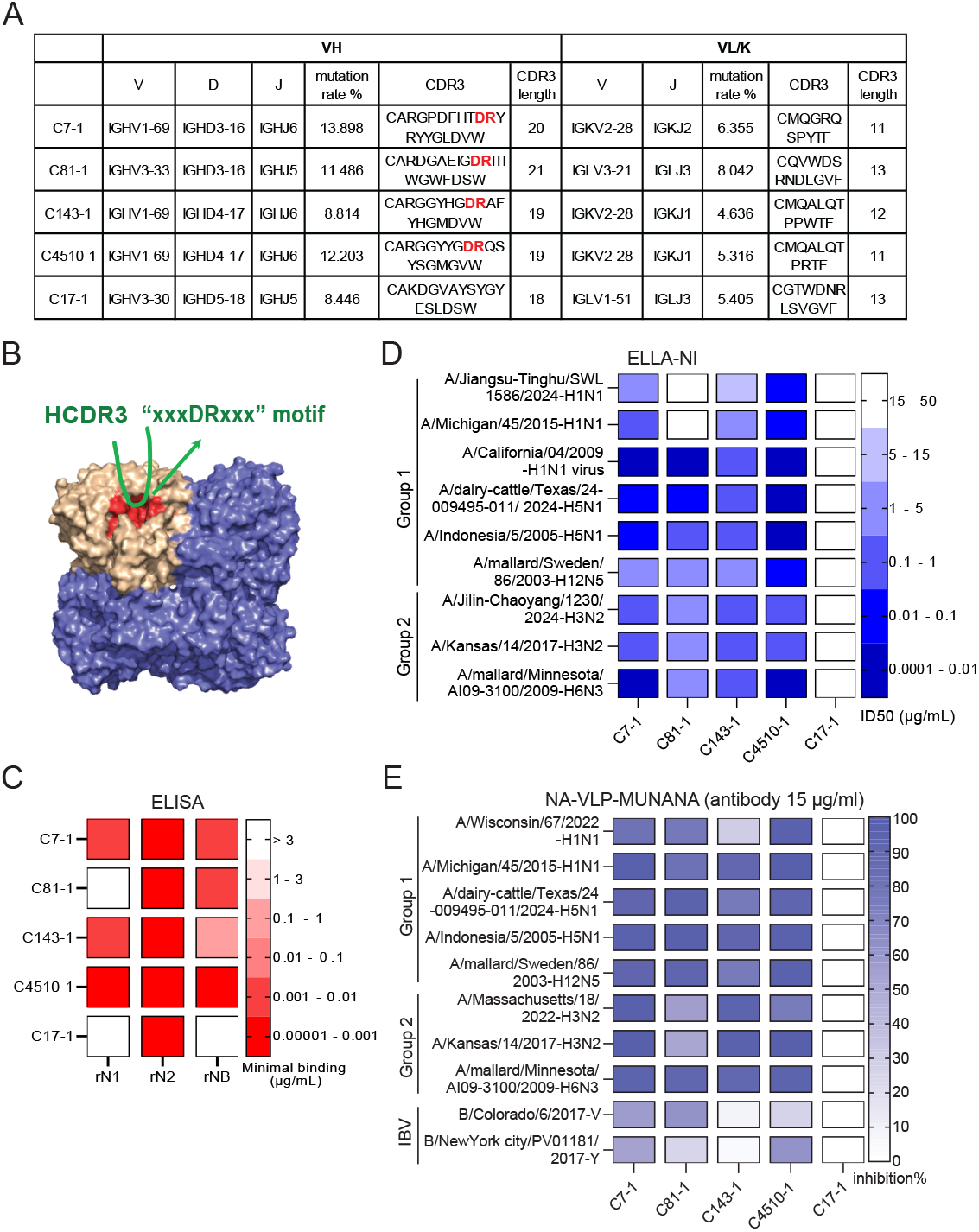
Broad NA-inhibition mAbs featuring an HCDR3 “xxxDRxxx” motif. (A) The usage of heavy and light chains genes, as well as the length and amino acid sequence of the HCDR3 for selected mAbs from the BCR repertoires of donor 3 (B) Schematic representation of the “DR” motif in the middle of HCDR3 interacts with NA conserved catalytic pocket. (C) Heat map of mAb binding to rN1(A/Michigan/45/2015-H1N1), rN2 (A/Kansas/14/2017-H3N2), and IBV-V rNB (B/Colorado/6/2017-Victoria lineage) measured by ELISA assay. The colored boxes represent the minimal binding concentration. (D) Heatmap of mAb inhibition activity measured by ELLA NI assay. The colored boxes indicate the half-maximal inhibitory concentration (infective dose [ID]50). The recombinant NA proteins from different influenza strains were utilized in the ELLA NI assay, except the A/California/04/2009-H1N1 viruses was applied. (E) Heatmap of mAb inhibition activity at 15 μg/mL against N1, N2, N5, N3, or IBV NA-VLPs measured by MUNANA assay. The colored boxes represent the percentage of inhibition.

We evaluated the binding of these mAbs to N1, N2, and NB by ELISA. C7-1, C143-1, and C4510-1 displayed cross-binding to N1, N2, and NB (Figure 3C). C81-1 showed cross-reactivity with N2 and NB, while C17-1 was specifically reactive to N2 (Figure 3C). To further evaluate the inhibition breadth of these mAbs, we established an NA panel derived from group 1 and group 2 for ELLA assay. We found that C7-1, C143-1, and C4510-1 showed broad inhibition activity against NAs from different influenza A viral strains. While C81-1 inhibited most NAs tested, no activity was detected for N1 (produced from insect cells) of A/Michigan/45/2015 and A/Jiangsu-Tinghu/SWL1586/2024 (Figure 3D). Since the ELLA assay uses a large-molecule substrate, the observed antibody inhibition may target the NA active site or surrounding epitopes. When small molecules are used as a substrate, as in the MUNANA-inhibition assay, only antibodies targeting the enzyme active site will exhibit inhibition. To further characterize the inhibition profile of the above mAbs, we performed a MUNANA assay using NA-viral-like particles (VLPs) generated by co-expressing HIV-1 gag, pol, and Rev with NA derived from IAV and IBV in HEK293T cells. The results demonstrated the broad inhibitory activity of C7-1, C143-1, C4510-1, and C81-1 against NA-VLPs from both IAV and IBV, including the recent cow H5N1 (Figure 3E). The inhibition differences of C81-1 against recombinant N1 (produced in insect cells) and N1-VLP (produced in 293T cells) are likely due to distinct N1 modifications, such as glycosylation, in insect and mammalian expression systems, which may prevent C81-1 recognition. Consistent with the ELLA results, antibody C17-1 did not exhibit any inhibition against any of the NA-VLPs tested in the MUNANA assay, suggesting that C17-1 recognized non-inhibitory epitopes of N2 (Figure 3E). Taken together, these findings emphasize that we can precisely locate NA BImAbs by using the described NA mAb signatures.

### HCDR3 “DR” motif localizes at the V-D or D-J junction and is essential for BImAb function

The rearrangement of V, D, and J segments plays a central role in sculpting the critical antigen-binding site in CDR3. Sequence analysis of the NA BImAbs described above revealed that the “DR” motif is predominantly located at the V-D or D-J junctional regions (Figure 4A). These BImAbs exhibited diverse D gene utilization. While the J gene did not directly encode “DR”, the J6 gene usage was most abundant (Figure 4A). To elucidate the structural basis of the interaction between “DR” and the NA active site, we modeled the complex structure of C7-1 with N1 derived from A/Jiangsu-Tinghu/SWL1586/2024 using AlphaFold. The structural model showed that “DR” is inserted into the conserved NA enzymatic pocket and interacts with two conserved NA R residues (Figure 4B). However, the actual interaction patterns between the “DR” motif and the NA active site require validation through structural experiments. To further test the function of the “DR” motif, we mutated “DR” to “AA” in C7-1 and C4510-1. ELLA results revealed that both C7-1_DR-AA and C4510-1_DR-AA mutants completely lost inhibition against N1 (A/Jiangsu-Tinghu/SWL1586/2024-H1N1) and N2 (A/Jilin-Chaoyang/1230/2024-H3N2) (Figure 4C). Notably, C7-1_DR-AA and C4510-1_DR-AA failed to inhibit the sialidase activity of N1-VLP (A/Wisconsin/67/2022), N2-VLP (A/Massachusetts/18/2022), and NB-VLP (B/Colorado/6/2017 and B/New York City/PV01181/2017) (Figure 4D). Together, these results demonstrate that the “DR” motifs serve as key sequence signatures for BImAbs to recognize diverse NAs.

**Figure 4.**
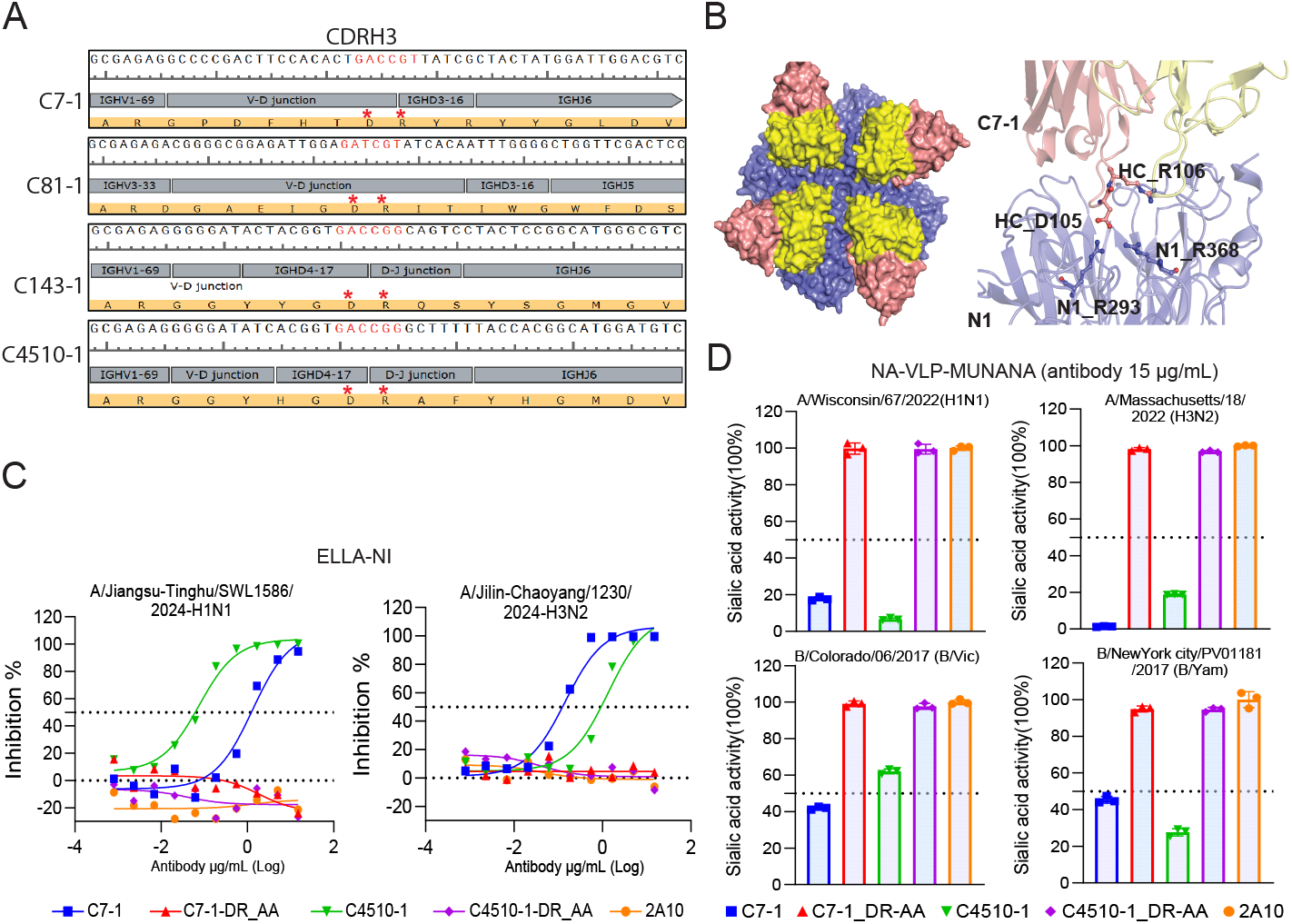
HCDR3 “DR” motif is crucial for BImAb function. (A) Schematic illustration of genetic characteristics of HCDR3 “DR” motif in BImAbs. The VDJ genes usage of CDRH3 sequence was characterized using the IgBlast (NCBI). The red asterisk indicates the location of the “DR” motif. (B) The structural prediction model of the binding of C7-1 (pink) to the NA enzymatic pocket (blue) using AlphaFold. N1 is derived from A/Jiangsu-Tinghu/SWL1586/2024. (C) Representative rN1 or rN2 inhibition curves by different wild-type and mutated NA mAbs in an ELLA assay. The dotted line represents 50% inhibition. (D) The wild-type and mutated mAb inhibition activity at 15 μg/mL against N1, N2, and IBV-V, or IBV-Y NA-VLPs measured by MUNANA assay.

### cMBC and aMBC-derived BImAbs exhibit similar NA inhibition activity

aMBC has been demonstrated to be associated with protection against malaria infection, and broadly neutralizing antibodies against Plasmodium falciparum have been identified from aMBC (28). However, it remains unclear whether aMBC serves as a source of broadly neutralizing or strain-specific mAbs against influenza. Here, we constructed the clonal evolutionary tree based on the VDJ gene sequences of the heavy chain derived from C7, C81 (BImAbs), and C17 (N2-specific binding mAb). We identified 15 clones for C7, 4 clones for C81, and 8 clones for C17, respectively (Figure 5A). These clones exhibited high and distinct V gene mutation rates compared to their germline V genes, indicating that these clonal B cells had undergone the germinal center response. We then utilized single-B cell transcriptome data to differentiate the MBC populations from which these clonal B cells originated. We found that C7-1, C7-10, C7-15, C81-3, C17-5, and C17-8 were located in the aMBC compartment, while all other clones exhibited cMBC signatures (Figure 5A). Both aMBC and cMBC-derived broad-reactive (C7 and C81) or N2-specific (C17) clones could be constructed under their germline clonal trees (Figure 5A). To further elucidate the function of aMBC-derived NA mAbs, we tested the inhibition activity of C7-1, C7-10, and C81-3 (aMBC-derived BImAbs) against H1N1 and H3N2 and compared it to the inhibition by C7-2, C7-5, and C81-1 (cMBC-derived BImAbs) (Figure 5B-C). We found that aMBC- and cMBC-derived BImAbs displayed comparable inhibition against H1N1 (2009) as tested by ELLA (Figure 5B, left panel). Consistently, the N1-VLP-based MUNANA assay demonstrated that these BImAbs exhibited similar inhibition against H1N1 (2022), with stronger inhibition by C7-10 and C81-3 being observed (Figure 5C, left panel). Furthermore, C7-1, C7-2, C7-5, and C7-10 similarly inhibited H3N2 in both the ELLA and MUNANA assays (Figure 5B-C, right panel). C81-1, which harbored a higher V gene mutation rate than C81-3, displayed balanced N1 and N2 inhibition, while C81-3 showed N1-biased inhibition (Figure 5C), suggesting clonal selection of C81 by N1 or N2 antigens. Additionally, C17-5 and C17-8 (aMBC-derived) exhibited similar binding to N2 compared to C17-1 and C17-2 (cMBC-derived) (Figure 5D). These data suggest that the development of NA broad-reactive MBCs may follow two distinct trajectories (cMBC and aMBC), but cMBC- and aMBC-derived BImAbs exhibit comparable functions.

**Figure 5.**
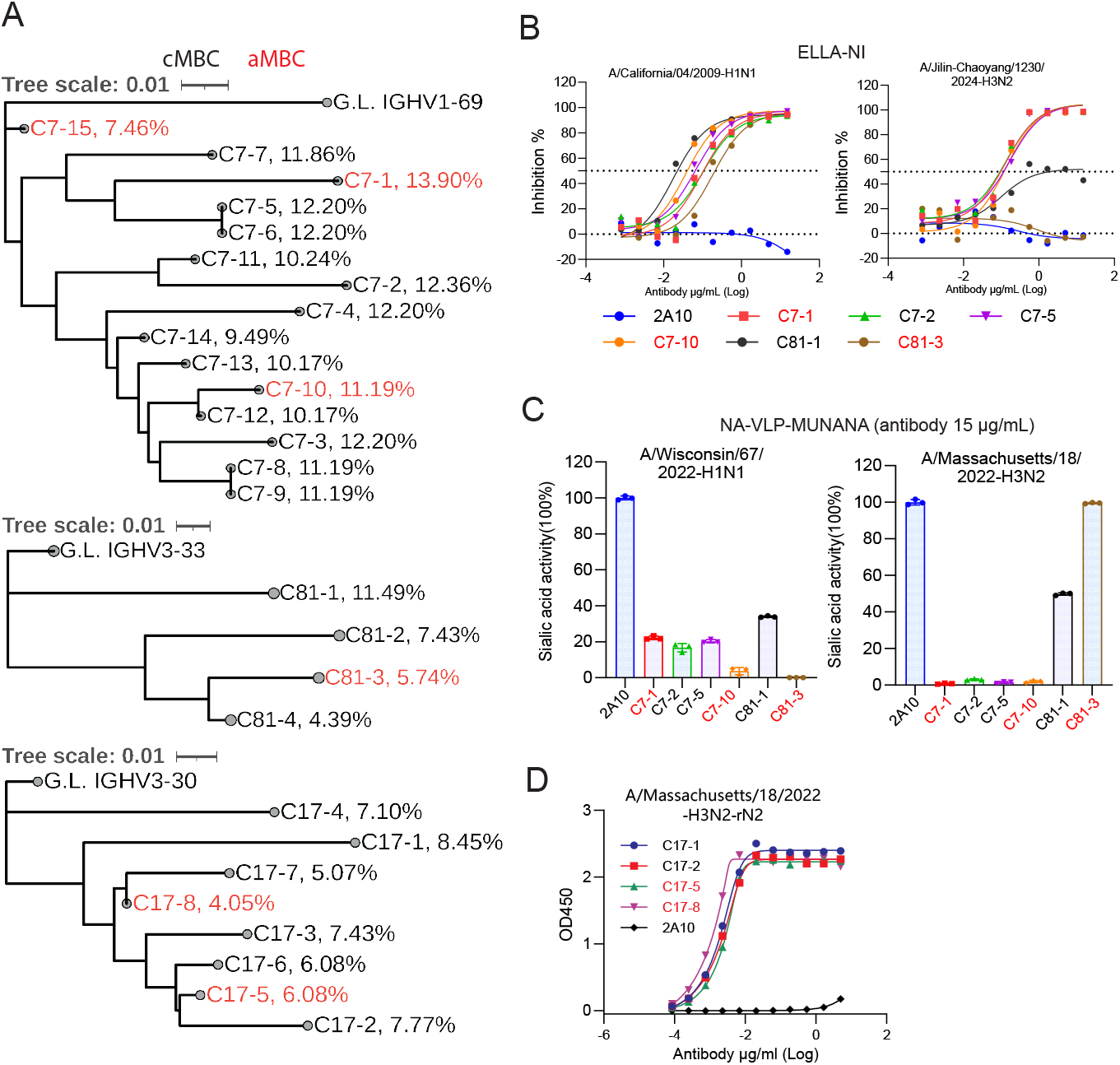
BImAbs derived from cMBC and aMBC exhibit NA inhibition activity. (A) The clonal evolutionary tree of the heavy chain VDJ sequence of C7, C81, or C17 clonotypes. Black and red indicate the B cell clones from cMBC and aMBC, respectively. The scale bar represents a 0.01% change in nucleotides. (B) Representative N1 (virus of A/California/04/2009-H1N1) or rN2 (recombinant NA protein from A/Jilin-Chaoyang/1230/2024-H3N2) inhibition curves by different NA mAbs in an ELLA assay. The dotted line represents 50% inhibition. (C) The mAb inhibition activity at 15 μg/mL against N1 and N2-VLPs measured by MUNANA assay. (D) The binding curves of different mAbs to rN2 (A/Jiangsu-Tinghu/SWL1586/2024-H3N2) measured by ELISA.

## Discussion

Influenza NA is the second most abundant protein on the viral surface and is essential for viral replication. There is growing interest in developing NA-based universal flu vaccines (35). However, our understanding of the molecular features, clonal selection, and developmental trajectories of NA broadly reactive B cells remains limited. This study identified a series of NA broadly reactive MBCs belonging to different clonal lineages and discovered that a long HCDR3 with an “xxxDRxxx” motif is an essential characteristic. Additionally, we found that NA broadly reactive MBCs follow a clonal evolutionary trajectory that spans both cMBC and aMBC compartments, which are two immunologically distinct populations. These findings advance our understanding of how NA broadly reactive B cells interact with NA-conserved protective epitopes and provide insights into the clonal evolution of these B cells. Targeting NA broadly reactive B cells may be an efficient approach for designing NA-based universal flu vaccines.

Consistent with previous studies (12, 14), antibodies inhibiting N1, N2, and IBV-V were detected in our cohort of 15 healthy individuals, indicating historical exposure to seasonal influenza. In our study, one out of 15 donors (donor 3) exhibited cross-inhibition titers against N5 and N3, while a previous study found one out of 200 healthy individuals (donor I) harbored high NA cross-reactive antibodies (17). The identification of such individuals provides confidence for the development of NA-based universal flu vaccines. One limitation of these studies is the lack of clarity regarding the influenza infection or vaccination history of the individuals, although such exposure history is likely associated with the levels of NA cross-reactive antibodies (36, 37). One recent study has shown that host genetics strongly drive biased influenza-subtype immune responses (38). Two individuals likely share common genetic features associated with the development of NA broadly reactive B cell responses. Indeed, the BImAb FNI19 (from a previous study) and C7-1, C143-1, and C4510-1 (isolated in this study) utilize the same VH1-69 germline gene, suggesting that VH1-69 usage in the B cell receptor (BCR) repertoire may be related to the NA broadly reactive B cell response. Interestingly, the VH1-69 germline gene is also highly utilized by hemagglutinin (HA) stem broadly neutralizing monoclonal antibodies (BnAbs) (39-42). The VH1-69 germline gene encodes the critical HCDR2 F54 residue, which is required for the initial development of stem BnAbs (42). In contrast, the VH1-69 NA BImAbs utilize a long HCDR3 with an “xxxDRxxx” motif to recognize the NA-conserved enzymatic pocket (12, 17). The elucidation of these molecular features of HA BnAbs and NA BImAbs advances our understanding of broad B cell responses against influenza virus.

In addition to VH1-69 NA BImAbs, we and others have identified NA BImAbs using VH3-30 (4N2C4), VH3-20 (1G01), VH3-33 (C81-1), and VH4-31 (DA03E17 and NCS.1) (12, 16-18, 43), indicating that the development of NA BImAbs is not restricted to VH1-69. One common feature of these NA BImAbs is a long HCDR3 with an “xxxDRxxx” motif. Structurally, the “DR” residues interact with the NA-conserved enzymatic pocket, mimicking sialic acid binding (12, 17, 43). Mutation of “DR” to “AA” fully abolishes the inhibition activity of NA BImAbs (Figure 4). Additionally, the “DR” residues are localized at the V-D or D-J junction, suggesting that the VDJ arrangement generating a long HCDR3 with an in-frame “xxxDRxxx” motif is the initial step in developing an NA broad B cell response. The frequency of broad precursor B cells determines B cell competitive fitness during the B cell response (44); thus, investigating the frequency of NA-specific B cells with these features using large BCR datasets will allow us to assess the feasibility of eliciting NA BImAbs response with NA-based vaccines. However, we cannot rule out the possibility that the “xxxDRxxx” motif is generated by SHMs during the NA-specific germinal center response. In this case, appropriate adjuvants or immunization strategies (45, 46) are likely required for NA vaccines to recruit NA-specific B cells into the germinal center to acquire “DR” mutations in the long HCDR3. The discovery of long HCDR3 with an “xxxDRxxx” motif for NA BImAbs not only guides the design of NA-based universal flu vaccines but also serves as a B cell immune marker to evaluate the efficiency of NA-based flu vaccines.

In the present study, we found that NA-specific MBCs from donor 3 were heterogeneous, consistent with our recent study using pooled NA-specific B cells (12). Two NA-specific MBC populations (aMBC and cMBC) were transcriptionally distinct, but the Ig isotype distribution and VH SHMs were comparable. (Figure 2). This immune phenotype is consistent with previously reported aMBCs in conditions such as malaria, HIV-1 infection, or systemic lupus erythematosus (SLE) (47, 48). Recent studies have demonstrated that aMBCs expand in response to malaria infection or influenza vaccination (26, 32, 49), and broadly neutralizing antibodies against *Plasmodium falciparum* have been cloned from aMBCs (24, 28). In our study, both aMBC- and cMBC-derived NA BImAbs exhibited similar NA inhibition functions, providing evidence that aMBCs form an essential MBC compartment for protection against influenza. The clonal lineage tree of BImAb C7 covered both cMBC and aMBC with high VH mutation rates, suggesting that naïve C7 precursor B cells were recruited into the germinal center by NA antigen, experienced clonal selection and then developed into cMBC and aMBC. However, the timing and mechanisms that regulate the formation of these two NA-specific MBC lineages require further investigation. In addition to identifying aMBC-derived NA BImAbs, we found aMBC-derived N2-specific monoclonal antibodies, indicating that antigen epitopes likely do not determine the fate-decision of aMBCs and cMBCs.

In conclusion, our study elucidated the molecular characteristics (long HCDR3 with an “xxxDRxxx” motif) of NA BImAbs, constructed the clonal evolutionary trajectory of NA BImAbs, and demonstrated that cMBCs and aMBCs are sources of NA BImAbs. These findings broaden our understanding of the development of NA BImAbs and provide a foundation for developing NA-based universal flu vaccines by targeting B cells with these features and stimulating them to mature and express NA BImAbs.

## Materials and Methods

### Human sample donors

Healthy individuals were recruited, and blood collections were performed by certified nurses to obtain plasma and PBMCs. The recruitment of healthy individuals was performed at the Center for Infectious Disease Research (CIDR) at Westlake University, and the protocols (20230613ZZL001 and 20231208ZZL001) were approved by the ethics committee of Westlake University. All participants signed the informed consent form, and the study conduction complied with all relevant ethical regulations. Compensations was provided to participants for their time and effort.

### Human PBMC and plasma isolation

Blood specimens were collected utilizing ethylenediaminetetraacetic acid (EDTA)-anticoagulated tubes. After collection, whole blood samples underwent centrifugation for 15 minutes at 1850 rpm. The plasma was then carefully removed from the cell pellet and stored at minus 80 °C. Peripheral blood mononuclear cells (PBMCs) were isolated by density-gradient sedimentation using Ficoll-Paque (Cytiva) as previously described (50). The isolated PBMCs were cryopreserved in heat-inactivated fetal bovine serum (FBS) supplemented with 10% dimethyl sulfoxide (DMSO) and subsequently stored in liquid nitrogen. Plasma samples were utilized for antibody quantification by enzyme-linked immunosorbent assay (ELISA) and enzyme-linked lectin assay (ELLA). The PBMC samples were used for single NA-specific B cell sorting.

### NA protein expression

The ectodomain head of the NAs from diverse influenza strains (Figure 3D) were cloned into a baculovirus transfer vector, pFastBac (Invitrogen) with an N-terminal gp67 signal peptide, 8xHis tag and the tetrabrachion tetramerization (TB) domain. The N2-R292K of A/H3N2/Kansas/14/2017 and NB-R292K of B/Colorado/6/2017 were also expressed as gp67-Avitag-8xHis tag-VASP-NA head and were used as NA probes. The constructed plasmid was transformed into DH10Bac competent cells to form a recombinant bacmid and the purified bacmid was transfected into Sf9 insect cells to produce recombinant baculovirus expression NA. To express the secreted NA protein, Hi5 cells were infected with recombinant baculovirus at MOI of 5 and incubated at 27 °C shaking at 150 RPM. When the viability of Hi5 cells reached 60%, the supernatant containing soluble NA proteins was harvested. The NA proteins were purified by His-Tag Protein Purification Resin (Beyotime). In some cases, the NA proteins were further purified by size exclusion chromatography on a Superose 6 or Superdex 200 (10/300GL) (Cytiva) column in 25 mM Tris pH8.0 with 150mM NaCl and 2mM CaCl_2_.

### Influenza NA ELISA

The 96-well half-area high-binding plates (Corning 3690) were coated with 50 μl per well of recombinant NA protein (2 μg/ml) diluted in PBS and incubated overnight at 4°C. For NA mAbs or polyclonal antibody detection, the plates were blocked with 3% bovine serum albumin (BSA) (BBI Life Sciences) in PBST for 1 hour at RT. Plates were washed 6 times with TBST (0.1% Tween-20). The mAbs or polyclonal antibodies were 3-fold serially diluted in PBST (0.05% Tween-20) with 1% BSA with a starting concentration of 5 μg/ml and 75 μl per well was added and incubated for 1.5 hours at RT. Plates were washed 6 times with TBST and 50 μl per well of Peroxidase-conjugated AffiniPure Goat anti-Human IgG (H+L) secondary antibody (Jackson) (1:20000) was added and incubated for 1.5 hours at RT. Plates were washed 6 times with TBST and 50 μl per well of TMB substrate (Thermo Fisher) was added. The plates were incubated for 5-7 min at RT and the color development was stopped by adding 50 μl per well of ELISA Stopping Solution (BBI Life Sciences). The absorbance was read at 450 nm with a microtiter plate reader and the data were analyzed with Microsoft Excel and GraphPad Prism.

### Influenza NA ELLA and ELLA-based inhibition assay

For ELLA-based titration of the recombinant NA or influenza virus, the flat-bottom 96-well MaxiSorp plates (Thermo Scientific) were coated for 16-20 hours at 4 °C with 150 μl/well of fetuin (Sigma) at 50 μg/ml. Fetuin-coated plates were blocked with 200 μl of 1% bovine BSA (BBI Life Sciences) in PBS for 1 hour at RT. The plates were washed 6 times with TBST (0.1% Tween-20), then the plates were incubated with 3-fold serial dilutions of recombinant NAs or influenza virus in PBS with 1 mM CaCl_2_ and 0.1% BSA for 16 hours at 37 °C. Following incubation, the plates were washed 6 times with TBST, then 100 μl of 2 μg/ml HRP-conjugated peanut agglutinin (Sigma) diluted in PBS was added. The plates were incubated for 1 hour and 45 mins at RT in the dark. After 6 times washing with TBST, 100 μl/well of TMB substrate (Thermo Fisher) was added. The plates were incubated for 5 mins at RT and the color development was stopped by adding 100 μl/well of ELISA Stopping Solution (BBI Life Sciences). The absorbance was read at 450 nm with a microtiter plate reader. The content of recombinant NAs or influenza virus that resulted in 90 to 95% of the maximum signal was chosen for use in the subsequent ELLA-based inhibition assay. To measure the NA inhibition (NI) titers, the mAbs were 3-fold serially diluted with a starting dilution of 15 μg/ml in PBS with 1 mM CaCl_2_ and mixed with the fixed amounts of recombinant NAs or influenza virus. Then, the mixtures were transferred to the fetuin-coated plates and incubated for 16 hours at 37 °C. Eight wells containing diluent without antibody were served as the positive (virus or NA-only) control. The flowing steps were the same as the above-described ELLA assay. Data were analyzed using Microsoft Excel and GraphPad Prism, and the 50% inhibitory dose (ID50) was defined as the concentration or dilution at which 50% of the NA activity was inhibited, compared to virus or NA-only control. Samples that did not reach 50% inhibition at the highly concentration of mAbs were considered as non-inhibition and the values were set to >15 μg/ml.

### Single B cell VDJ and transcriptome sequencing

B cells were isolated from frozen PBMCs of donor 3 by immunomagnetic negative selection according to the manufacturer’s protocol (EasySep Human B Cell Enrichment Kit, STEMCELL). The purified B cells were first stained with 1200 ng N2-M5 probes (600 ng per color) for 30 min at 4 °C, then stained with CD19-BV605 (Biolegend, 363023), CD27-BB515 (BD, 564643), IgD-AF700 (Biolegend, 348230), and 7-AAD (Biolegend, 420403). The live N2-specific MBCs were sorted into the FCS buffer. The live N2 negative MBCs were sorted and stained with NB-M5 probes. The NB-specific MBCs were sorted and pooled with N2-specific MBCs, and directly loaded on a 10X Chromium A chip. Single-cell lysis and cDNA synthesis were performed by using the 5’ library and gel bead kit according to the provided protocol. The first-strand cDNA of 10X Barcoded was purified by silane magnetic beads, and sufficient mass was obtained by PCR amplification. The PCR products were quantified and qualified by Qubit3.0 Fluorometer (Invitrogen) and Agilent 2100 Bioanalyzer (Agilent). Then, the transcriptome and VDJ library preparation is performed by using 10X Chromium Single Cell VDJ Kits. All libraries were quantified by using Qsep100 (Bioptic) and Qubit3.0 Fluorometer (Invitrogen). Then libraries with different indices were multiplexed and loaded on an Illumina NovaSeq instrument according to the manufacturer’s instructions (Illumina). Sequencing was carried out using a 2x150bp paired-end (PE) configuration; image analysis and base calling were conducted by the NovaSeq Control Software (NCS) + OLB + GAPipeline-1.6 (Illumina) on the NovaSeq instrument.

### Single-B cell RNA-seq and VDJ-seq data analysis

The raw fastq files were processed using the 10X Genomics CellRanger (7.1.0) pipeline. The cell quality control, clustering, and marker gene analysis were performed by using Seurat (v4.3.0). Genes expressed in over 10 cells were retained, and cells were filtered based on the number of genes and the mitochondrial gene percentage to remove possible doublets. Cell types were identified by using SingleR. B cell VDJ contig assembly, annotation, and clonotype analysis were performed using ‘‘cellranger vdj’’ with the Cell Ranger V(D)J compatible reference. The somatic hypermutation rate was calculated by mapping VDJ sequences against the germline reference from the international ImMunoGeneTics information system (IMGT) or IgBlast (NCBI).

### Monoclonal antibody cloning and expression

The heavy and light chain VDJ fragments were cloned into IgG1 and IgK or IgL expression vectors. The heavy and light chain plasmids were co-transfected into Expi293F cells at a ratio of 1:1 and antibodies were purified from supernatant using Protein A agarose resin. The purified antibodies were further buffer exchanged into PBS and concentrated. The antibodies were sterilized by 0.22 μm filtration.

### Generation of NA-VLP

To generate NA-Viral Like Particles (VLP), HEK293T cells were co-transfected with 750 ng of pMDLg/pRRE, 250 ng of pRSV-Rev, and 800 ng of pcDNA3.1 encoding the indicated NA (Figure 3E) by using Lipofectamine 3000 (Thermo) according to the manufacturer’s instructions. After 2 days, the supernatant of the transfected cells was harvested and stored at minus 80°C in aliquots.

### NA-MUNANA assay

The NA-Fluor™ Influenza Neuraminidase Assay Kit (Applied Biosystems) was employed to test the NA-inhibition activities of mAbs. Briefly, mAbs (15 μg/ml) were mixed with a fixed amount of NA-VLP in 96-well black plates (Thermo) and incubated for 30 min at 37°C. Subsequently, the NA-Fluor™ substrate (MUNANA) was added at a final concentration of 200 μM. The plates were incubated for 1 hour at 37 °C, then the reaction was stopped with NA-Fluor™ stop solution and the NA activity was measured by fluorescence using Synergy H1 microplate reader (BioTek) (excitation at 365 nm and emission at 445 nm).

### Construction of NA mAbs clonal evolutionary tree

For germline gene reconstruction, the CDR3 loops of germline sequences were replaced with the corresponding region from the B cell clone with the lowest VH mutation rate within each clonotype, as it was not possible to unambiguously determine the original identity of the junction region. The heavy chain VDJ sequences were aligned, and phylogenetic trees were subsequently constructed in MEGA (version 11) using the neighbor-joining algorithm, with rooting based on the closest germline immunoglobulin gene matches. These phylogenetic trees were then annotated and visualized through the iTOL platform.

### Software used for data analysis

Data and statistical analyses were performed in FlowJo 10 and GraphPad Prism unless otherwise stated. The complex structure of C7-1 with N1 of A/Jiangsu-Tinghu/SWL1586 was modeled by the Alpha fold server and visualized in Pymol. Phylogenetic trees were performed in MEGA (version 11) and visualized in iTOL platform.

## Acknowledgements

We thank all the donors for their blood donation. We thank the clinical physician for the human blood collection. We thank the flow cytometry core at Westlake University for assistance in B cell sorting. This project is supported by the Westlake Educational Foundation to Z.Z., National Natural Science Foundation of China (NSFC) to Z.Z. (82471855).

## Author contributions

Conceptualization: Z.Z.; Methodology: C.L., Q.Y.; Formal analysis: C.L., Q.Y.; Investigation: C.L., Q.Y.; Donor recruitment: L.H., X.W.; Funding acquisition: Z.Z.; Writing: C.L., Z.Z.; Supervision: Z.Z.

